# SLC17 transporters mediate renal excretion of Lac-Phe in mice and humans

**DOI:** 10.1101/2024.04.18.589815

**Authors:** Veronica L. Li, Shuke Xiao, Pascal Schlosser, Nora Scherer, Amanda L. Wiggenhorn, Jan Spaas, Alan Sheng-Hwa Tung, Edward D. Karoly, Anna Köttgen, Jonathan Z. Long

**Affiliations:** Department of Pathology, Stanford University School of Medicine, Stanford, CA, USA; Department of Chemistry, Stanford University, Stanford, CA, USA; Sarafan ChEM-H, Stanford University, Stanford, CA, USA; Stanford Cardiovascular Institute, Stanford University, Stanford, CA, USA; Stanford Diabetes Research Center, Stanford University, Stanford, CA, USA; The Phil & Penny Knight Initiative for Brain Resilience at the Wu Tsai Neurosciences Institute, Stanford University, Stanford, CA, USA; Institute of Genetic Epidemiology, Faculty of Medicine and Medical Center, University of Freiburg, Freiburg Germany; Department of Epidemiology, Johns Hopkins University Bloomberg School of Public Health, Baltimore, Maryland, USA; Spemann Graduate School of Biology and Medicine (SGBM), University of Freiburg, Freiburg, Germany; Metabolon, Inc.

## Abstract

N-lactoyl-phenylalanine (Lac-Phe) is a lactate-derived metabolite that suppresses food intake and body weight. Little is known about the mechanisms that mediate Lac-Phe transport across cell membranes. Here we identify SLC17A1 and SLC17A3, two kidney-restricted plasma membrane-localized solute carriers, as physiologic urine Lac-Phe transporters. In cell culture, SLC17A1/3 exhibit high Lac-Phe efflux activity. In humans, levels of Lac-Phe in urine exhibit a strong genetic association with the *SLC17A1-4* locus. Urine Lac-Phe levels are also increased following a Wingate sprint test. In mice, genetic ablation of either SLC17A1 or SLC17A3 reduces urine Lac-Phe levels. Despite these differences, both knockout strains have normal blood Lac-Phe and body weights, demonstrating that urine and plasma Lac-Phe pools are functionally and biochemically de-coupled. Together, these data establish SLC17 family members as the physiologic urine transporters for Lac-Phe and uncover a biochemical pathway for the renal excretion of this signaling metabolite.

## Introduction

Lactate is a fundamental metabolite that has attracted significant biochemical and physiologic attention for over a century (1, 2). Classically, lactate is produced from glucose when energy demands are high, such as in muscle during exercise, and can reach millimolar concentrations in the circulation (3, 4). Once produced, lactate exerts pleotropic functions in systemic energy metabolism. These include serving as a metabolic fuel, tissue redox shuttling, signaling via cell surface receptors, and modification of proteins via lysine lactoylation (5–8).

Recently, we showed that the metabolic functions of lactate also extend to lactate-derived metabolites. Specifically, we identified N-lactoyl-phenylalanine (Lac-Phe) as an exercise- and metformin-inducible signaling molecule that regulates food intake and energy balance (9–11). Lac-Phe is produced via the action of a cytosolic enzyme called CNDP2 that catalyzes the direct condensation of lactate and phenylalanine (12). Genetic ablation of CNDP2 in mice resulted in dramatic reduction in circulating basal and stimulated Lac-Phe levels. CNDP2-KO mice also exhibited a hyperphagic and obese phenotype after concurrent exercise training and high fat diet feeding (9), establishing that Lac-Phe is necessary for the suppression of feeding after exercise training. CNDP2-KO mice are also resistant to the anorexigenic and body weight-lowering effects of metformin therapy (10), a pharmacological stimulus that increases glycolysis and lactate flux. Conversely, pharmacological administration of Lac-Phe is sufficient to suppress feeding and obesity without affecting energy expenditure. In humans, higher post-exercise Lac-Phe levels are associated with greater reduction of adiposity following exercise training (13), suggesting that the Lac-Phe pathway is conserved across multiple mammalian species.

The molecular mechanisms by which Lac-Phe is transported from the cytosol to the extracellular space has remained elusive. Three lines of evidence point to the existence of a specific transporter that mediates active Lac-Phe efflux from cells. First, CNDP2 is an intracellular, cytosolic enzyme, yet Lac-Phe is enriched in the conditioned medium of cultured cells (9). This intracellular versus extracellular distribution of CNDP2 and Lac-Phe, respectively, suggest active processes mediate efflux of Lac-Phe away from its site of production. Second, Lac-Phe is a polar metabolite and thus would not be expected to simply diffuse across lipid bilayers. Third, Lac-Phe can be actively translocated across the plasma membrane by transporters in cell culture. For instance, overexpression of ABCC5, one member of the ABC transporter family, increases Lac-Phe levels in conditioned media of cells (12). However, there is no evidence that Lac-Phe levels are altered in ABCC5-KO mice (9), suggesting that other, as of yet unidentified plasma membrane carriers are the physiologically relevant transporters for Lac-Phe *in vivo*.

Here we combine human metabolite-genome association data with biochemical studies *in vitro* and genetic studies in mice to demonstrate that Lac-Phe is an endogenous transport substrate for the SLC17 family of solute carriers. SLC17A1/3 are robust effluxers of Lac-Phe in cell culture and highly expressed in renal epithelial cells in mice and humans. Genetic individual ablation of SLC17A1 or SLC17A3 in mice results in reduced urine levels of Lac-Phe. Finally, we show that both SLC17A1- and SLC17A3-KO mice have normal levels of Lac-Phe in blood and normal body weights, providing evidence for the existence of yet additional transporters that mediate Lac-Phe efflux into different biological fluids.

## Results

### Re-assignment of 1-carboxyethylphenylalanine as Lac-Phe and its genetic association with the SLC17A1-4 locus in human urine

In a 2020 metabolite genome-wide association study (14), we noted the detection of a urine metabolite which was reported by Metabolon as both “X-15497” and also “X-15497-retired for 1-carboxyethylphenylalanine”. In a subsequent 2023 study from the same group (15), this X-15497/1-carboxyethylphenylalanine metabolite was re-assigned by Metabolon to a different chemical structure, N-lactoyl-phenylalanine (Lac-Phe). In both studies, urine levels of the X-15497/1-carboxyethylphenylalanine/Lac-Phe metabolite exhibited a strong genetic association with the human *SLC17A1-4* locus (**Fig. 1a-c**). While 1-carboxyethylphenylalanine and Lac-Phe have identical molecular formulas and masses (C_12_H_15_NO_4_, [M-H]^-^ = 236.0928), these two molecules are in fact two distinct chemical entities. Until now, raw chromatographic or tandem mass spectrometry data were not available in either the 2020 or 2023 studies. Consequently, the precise chemical structure of this X-15497/1-carboxyethylphenylalanine/Lac-Phe metabolite, and the functional significance of its association with the *SLC17A1-4* locus, remained undefined. We first sought to definitively determine the chemical structure of the X-15497/1-carboxyethylphenylalanine/Lac-Phe metabolite. First, we compared tandem mass spectrometry (MS/MS) spectra of authentic standards for Lac-Phe and 1-carboxyethylphenylalanine. Both exhibited a major daughter ion of m/z = 88 corresponding to fragmentation of the amide N-Calpha bond (**Fig. 1e**). In addition, the 1-carboxyethylphenylalanine standard produced a minor daughter ion of m/z = 192 which was absent in the Lac-Phe standard; we presume this m/z = 192 daughter ion likely corresponds to decarboxylation of the 1-carboxyethyl portion of the molecule (**Fig. 1e**). The experimental MS/MS for X-15497, which was obtained from Metabolon, also showed the major m/z = 88 and no evidence for the m/z = 192 daughter ion. These data demonstrate that the fragmentation pattern for the metabolite X-15497 is more alike to Lac-Phe than it is to 1-carboxyethylphenylalanine (**Fig. 1e**).

**Fig. 1.**
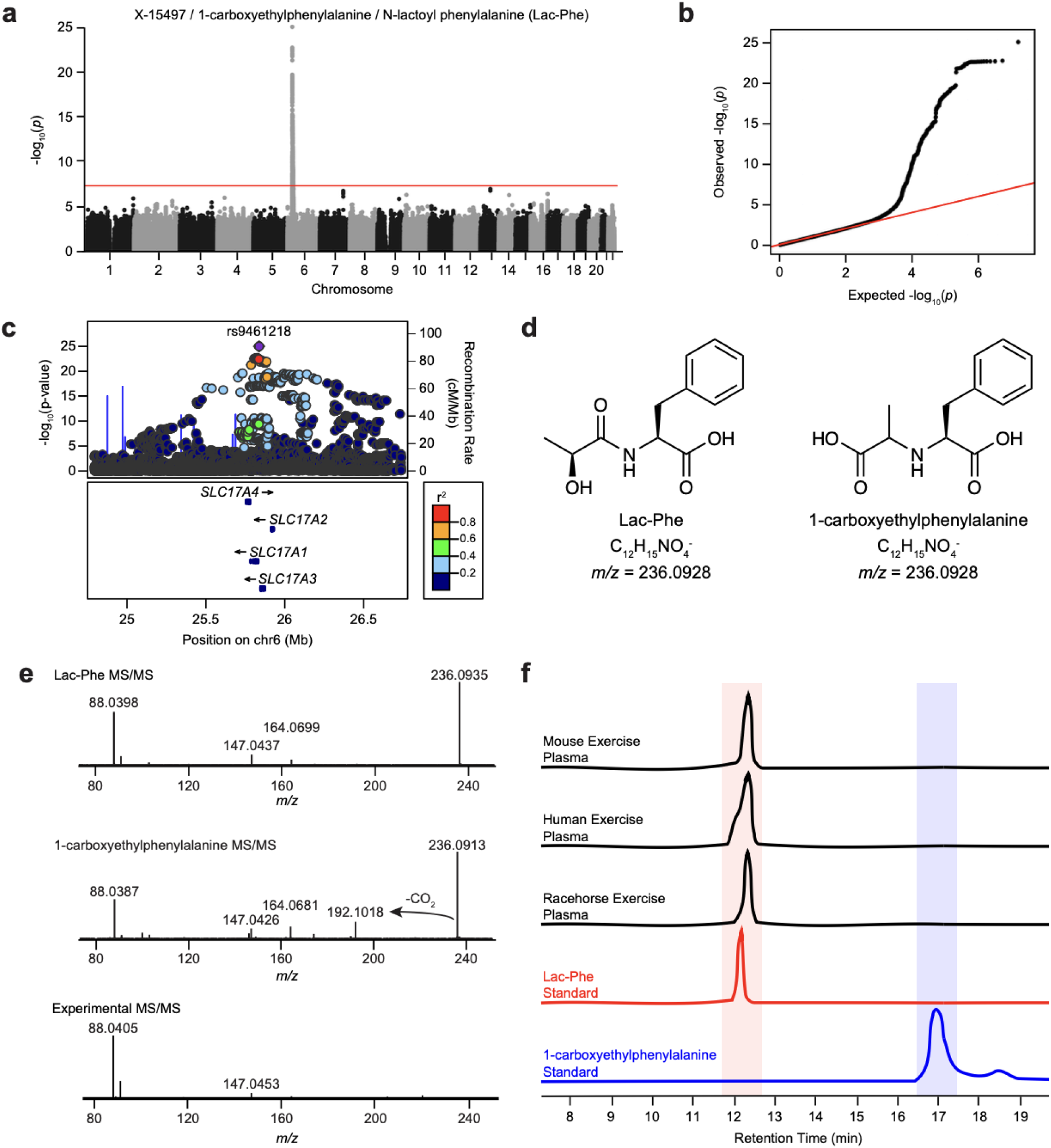
Structural assignment and association of X-15497/1-carboxyethylphenylalanine/Lac-Phe with the *SLC17A1-4* locus in humans. (**a-c**) Manhattan (**a**), QQ (**b**), and regional association plot (**c**) of the genetic association of the urine metabolite, X-15497/1-carboxyethylphenylalanine/N-lactoyl-phenylalanine (Lac-Phe), to the *SLC17A1-4* locus on chromosome 6. (**d**) Chemical structure of Lac-Phe (left) and 1-carboxyethylphenylalanine (right). (**e**) Tandem mass spectrometry fragmentation of an authentic Lac-Phe standard (top), an authentic 1-carboxyethylphenylalanine standard (middle), and the experimental X-15497/1-carboxyethylphenylalanine/N-lactoyl-phenylalanine metabolite (bottom). (**f**) Extracted ion chromatograms of the endogenous 236.0928 peak in mouse, human, and racehorse plasma along with synthetic Lac-Phe and 1-carboxyethylphe standards.

We also compared the retention time for the endogenous m/z = 236.0928 peak from blood plasma and authentic samples of Lac-Phe and 1-carboxyethylphenylalanine standards by liquid chromatography-mass spectrometry (LC-MS). As shown in **Fig. 1f**, in mouse, human, and racehorse plasma, we detected a m/z = 236.0928 peak eluting at ∼12 min, which was an identical retention time to that of the Lac-Phe standard. By contrast, 1-carboxyethylphenylalanine eluted five minutes later at ∼17 min. In fact, no endogenous peak at m/z = 236.0928 and a 17 min retention time could be observed. On the basis of both MS/MS fragmentation and retention time, we conclude that the X-15497/1-carboxyethylphenylalanine/Lac-Phe metabolite is indeed Lac-Phe, and not 1-carboxyethylphenylalnine. In addition, our data demonstrate that 1-carboxyethylphenylalanine is not an endogenous metabolite.

### SLC17 family members can mediate Lac-Phe efflux in vitro

Having confirmed the chemical structure for X-15497/1-carboxyethylphenylalanine as Lac-Phe, we next turned to the observed genetic association of urine Lac-Phe with the *SLC17A1-4* locus. The lead single nucleotide polymorphism (SNP), rs9461218, is located in an intronic region of the *SLC17A1* gene and is associated with a decrease in urine Lac-Phe levels (beta = -0.242 +/- 0.023, P = 9.9e-26, non-effect allele G, effect allele A, effect allele frequency 0.22) (15). However, the genetic association signal maps into an extended linkage disequilibrium block and encompasses the adjacent genes *SLC17A2*, *SLC17A3*, and *SLC17A4* (**Fig. 1c**). No genetic association signal was detected between Lac-Phe levels in blood and the *SLC17A1-4* locus (15). A significant correlation was detected between urine and plasma Lac-Phe levels (r = 0.54, P < 2.2e-16, **Fig. S1**). In the Type 2 Diabetes Knowledge Portal (T2D-KP), the *SLC17A1-4* locus is significantly associated with multiple cardiometabolic traits (**Fig. S2**).

The *SLC17A1-4* genes encode for four plasma membrane solute carriers, which are also called NPT1-4. SLC17A1-4 transporters were originally shown to be co-transporters of sodium and phosphate *in vitro*. Other additional *in vitro* transport substrates include urate and synthetic organic anions such as p-aminohippuric acid (16–18). Moreover, human genetic screens of entire plasma and urine metabolomes point towards dozens of putative additional physiological and xenobiotic substrates (15). A biochemical interpretation of the genetic association between Lac-Phe and the *SLC17A1-4* locus is that Lac-Phe is an endogenous substrate transported by SLC17 family members. To test this prediction, we obtained mammalian expression plasmids for each SLC17A1-4 and cDNAs were individually transfected into HEK293T cells. As an additional positive control, we also transfected ABCC5, a previously reported in vitro Lac-Phe transporter (12). One day after transfection, cells were changed to serum free media and incubated for an additional 24 h. Media Lac-Phe levels were then measured by LC-MS (**Fig. 2a**). Compared to GFP-transfected controls, we observed a 2-fold and 5-fold increase in extracellular Lac-Phe in cells transfected with SLC17A1 and SLC17A3, respectively (**Fig. 2b**). By contrast, cells transfected with SLC17A2 and SLC17A4 did not exhibit any significant increase in media Lac-Phe levels. ABCC5-tranfected cells also exhibited a ∼2-fold increase in Lac-Phe. The mRNA expression for each SLC17A1-4 was comparable as measured by qPCR (**Fig. S3a**). Western blotting of transfected lysates with an anti-flag antibody confirmed protein expression of SLC17A2-4, while protein levels of SLC17A1 were below the limit of detection (**Fig. S3b**). We conclude that overexpression of SLC17A3, and to a lesser extent SLC17A1, is sufficient to drive Lac-Phe efflux from cells.

**Fig. 2.**
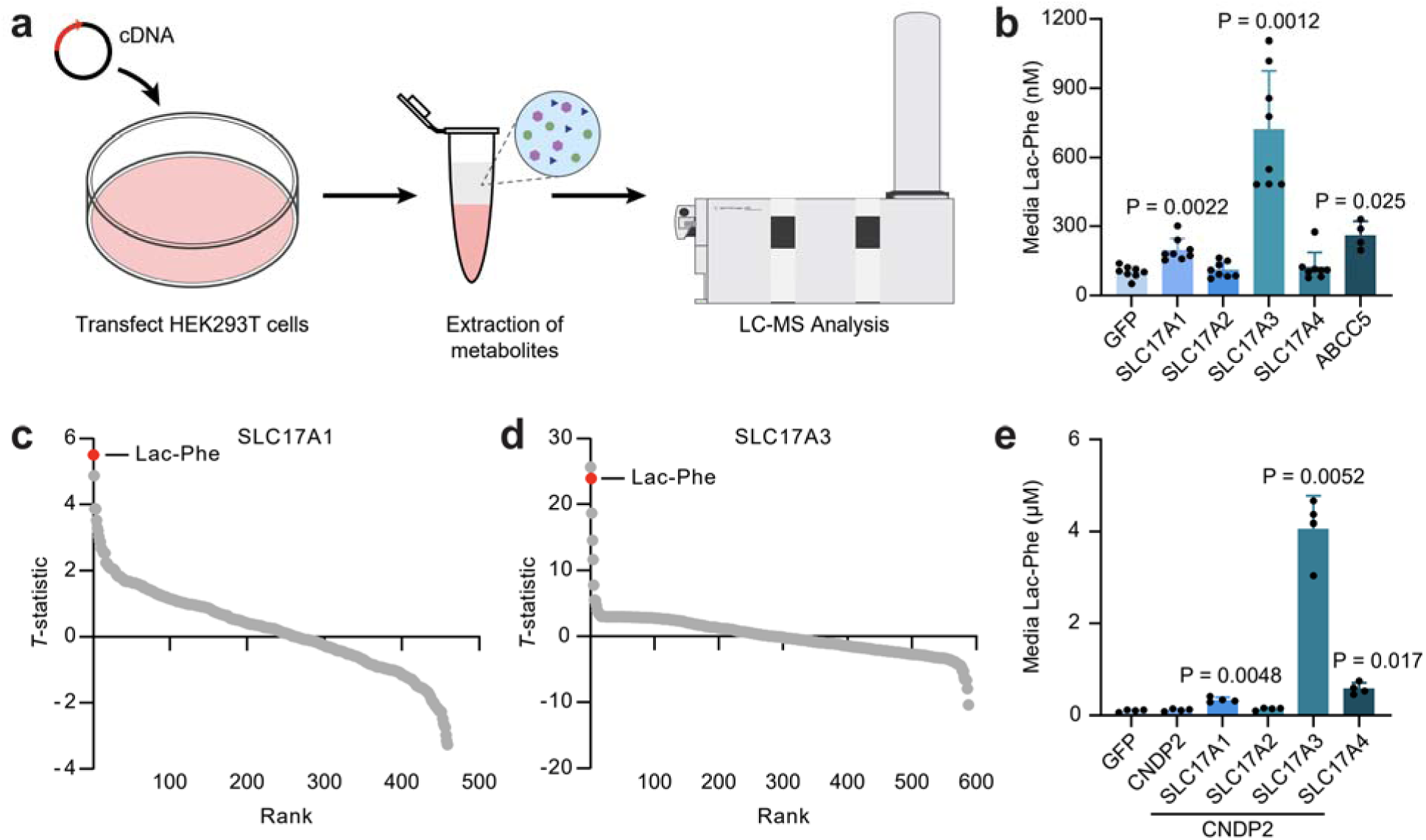
Overexpression of SLC17 family members drives Lac-Phe secretion *in vitro*. (**a**) Schematic of *in vitro* secretion assay. (**b**) Lac-Phe levels in conditioned medium of GFP or SLC17A1-4 transfected cells. (**c,d**) T-statistic of all conditioned media peaks detected by untargeted metabolomics in GFP versus SLC17A1 (**c**) or SLC17A3 (**d**) transfected cells. (**e**) Lac-Phe levels in conditioned medium of cells co-transfected with CNDP2 and the indicated SLC17 family member or just with GFP as a control. For **b**, N=8/group, except ABCC5 where N=4/group. For **c-e**, N=4/group. Data are shown as mean ± SD. P-values in (b) and (e) were calculated by one-way ANOVA and Dunnett T3 post-hoc tests.

Next, we performed untargeted metabolomic comparisons of conditioned media from SLC17A1-transfected or SLC17A3-transfected cells compared to GFP controls. Using XCMS software (19), we identified a total of N=13 and N=20 features that were significantly increased (P<0.05) upon SLC17A1 or SLC17A3 transfection, respectively, compared to GFP control (**Table S1** and **Fig. 2c,d**). Notably, Lac-Phe was the top most increased metabolite in SLC17A1-transfected media, and it was the second most increased metabolite in SLC17A3-transfected media (**Fig. 2c,d**). These data demonstrate that amongst all detected metabolites, Lac-Phe is a principal transport substrate for SLC17A1 and SLC17A3.

If SLC17 transporters mediate efflux of Lac-Phe, then overexpression of both CNDP2 and SLC17 transporters should result in even greater levels of extracellular Lac-Phe because now both intracellular biosynthesis and plasma membrane efflux are increased. We therefore transfected CNDP2 alone or co-transfected CNDP2 with each individual SLC17 transporter. CNDP2 overexpression was confirmed by Western blot (**Fig. S3c**). Overexpression of CNDP2 alone did not alter media Lac-Phe levels, establishing that driving intracellular biosynthesis alone is not sufficient to increase extracellular Lac-Phe levels. However, when CNDP2 was overexpressed with SLC17A1 or SLC17A3, we observed a dramatic 3-fold and 40-fold increase in extracellular Lac-Phe levels, respectively (**Fig. 2e**). We also observed a synergistic effect of overexpression of CNDP2 and SLC17A4: in this case, media Lac-Phe levels were increased by 6-fold, whereas no changes were observed with CNDP2 or SLC17A4 overexpression alone. Lastly, no increase in media Lac-Phe could be observed with SLC17A2. Quantification of intracellular levels of phenylalanine and Lac-Phe did not reveal any major differences upon transfection by individual SLC17 family members alone or in combination with CNDP2; intracellular lactate was below the limit of detection of our method (**Fig. S3d-i**). We also did not observe any evidence for Lac-Phe accumulating intracellularly upon CNDP2 overexpression alone, which potentially reflects alternative pathways of intracellular Lac-Phe metabolism (**Fig. S3d-i**). These data show that CNDP2 and SLC17 family members can cooperatively increase extracellular Lac-Phe levels.

To determine the generality of SLC17-mediated Lac-Phe efflux in non-renal cells, we also measured Lac-Phe levels in media after transfection of the mouse macrophage cell line RAW264.7. Overexpression of SLC17A2 and SLC17A3, but not SLC17A1 or SLC17A4, in RAW264.7 macrophages increased media Lac-Phe levels by ∼50% (**Fig. S4a**). Therefore, the magnitude of SLC17-mediated Lac-Phe efflux in cell culture is dependent both on the specific transporter and also the cell type.

We used cellular knockout studies to determine whether genetic ablation of SLC17 family members would reduce Lac-Phe efflux in vitro. HEK293T cells do not express SLC17A proteins (**Fig. S3a**, Ct > 30). We therefore turned to the mouse kidney epithelial TKPTS cells. By mRNA levels, only *Slc17a1* was robustly expressed in this cell line while mRNA levels for *Slc17a2*, *Slc17a3*, and *Slc17a4* were not detected (**Fig. S4b**). In SLC17A1-KO TKPTS cells, media Lac-Phe levels were reduced by 40% compared to control TKPTS cells (**Fig. S4c**), demonstrating that SLC17 family members are physiologic contributors to Lac-Phe efflux in vitro.

### Urine Lac-Phe levels after sprint exercise

Next, we examined the tissue mRNA expression pattern for SLC17A1-4 in both mice (using qPCR analysis of a tissue panel) and humans (using GTEx) (20). In both species, SLC17A1/3, the two transporters with greatest cellular Lac-Phe efflux activities, were highly expressed in the kidney (**Fig. 3a,b** and **Fig. S5**). On the other hand, human *SLC17A2* was liver-specific and human *SLC17A4* exhibited a more broad tissue expression pattern (**Fig. S5**). The public mouse gene expression portal BioGPS (21) also confirmed restricted expression of *Slc17a1/3* to the kidney (**Fig. S6**). Lastly, within the kidney, the mouse single cell RNAseq dataset Tabula Muris showed high enrichment of *Slc17a1/3* in epithelial cells of the proximal tubule, which, coincidentally, was also where high expression of *Cndp2* mRNA was found (**Fig. S7**). These expression data suggest that SLC17A1/3-dependent secretion of Lac-Phe may be most relevant for regulating urine levels of Lac-Phe.

**Fig. 3.**
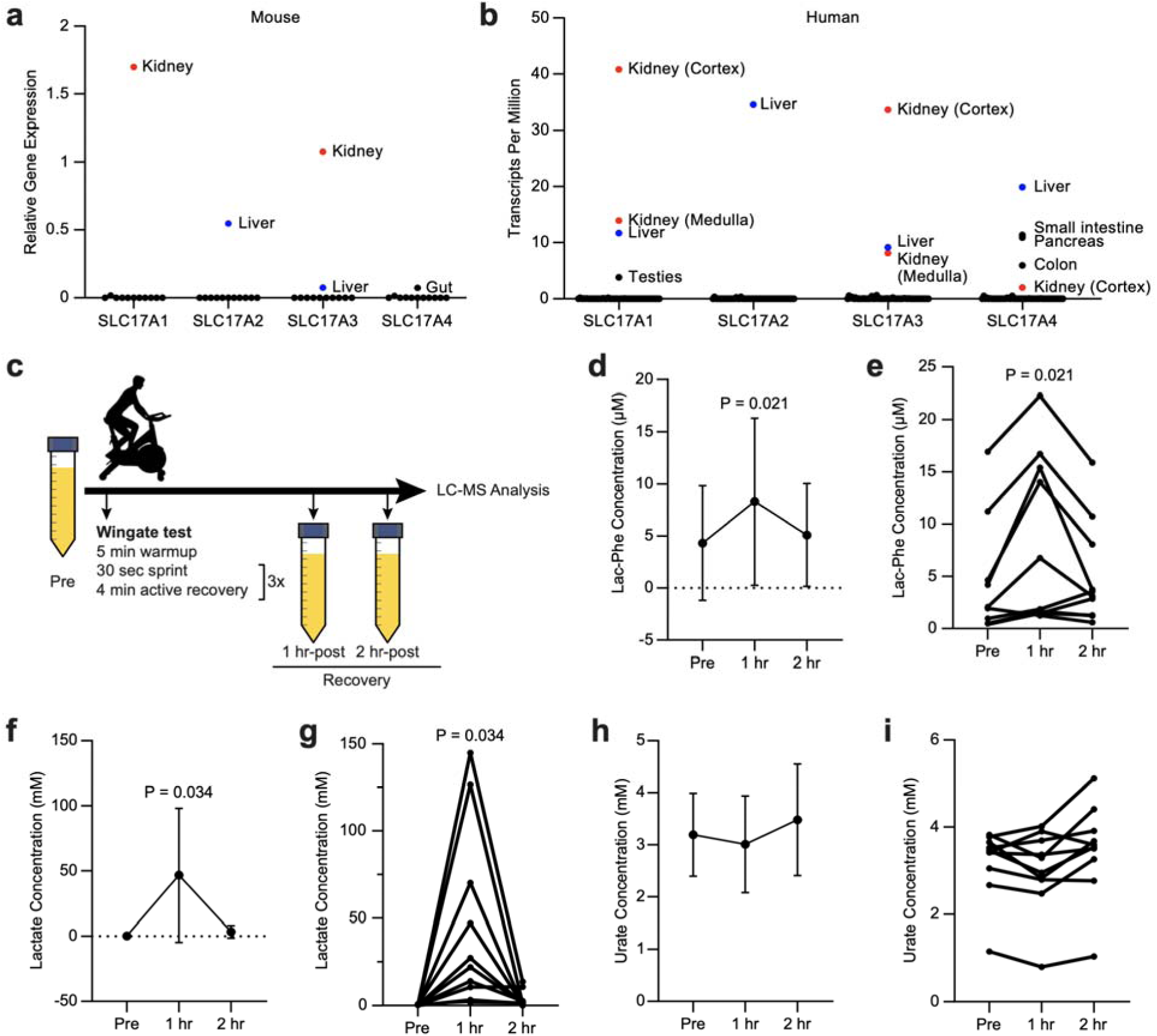
Lac-Phe levels increase in human urine after sprint exercise. (**a,b**) mRNA expression levels of SLC17 family members across mouse (**a**) and human (**b**) tissues. Mouse data were obtained from bioGPS; human data from GTEx. (**c**) Schematic of human sprint exercise study design. (**d-i**) Average (**d,f,h**) and individual (**e,g,i**) urine levels of the indicated metabolite before (pre) or at the indicated time point after exercise. For **d-i**, subjects were 6 male, 4 female, age 28.3 ± 5.3 (mean ± SD), N=10/group. Data are shown as mean ± SD. P-values in (**d-i**) were calculated by one-way ANOVA with Dunnett T3 post-hoc tests.

To determine if urine Lac-Phe levels are dynamically regulated by exercise, we measured Lac-Phe before and after a Wingate sprint test in humans. In our previous Wingate test study (9), urine was not collected. We therefore performed a new Wingate test in N=10 healthy volunteers and collected urine before and at 1 and 2 h after the test (**Fig. 3c**). This Wingate test consisted of three sets of sprinting (30 s each sprint) on a bicycle ergometer. We observed a wide range of basal urine Lac-Phe concentrations in humans (range: 0.5-17 uM, average 4 uM). One hour after the sprint exercise, urine Lac-Phe levels were increased by 2-fold and returned back to baseline by 2 h post-exercise (**Fig. 3d,e**). In addition to Lac-Phe, we also measured urine levels of both lactate and urate. Urine lactate also exhibited similar kinetics to Lac-Phe, being dramatically induced by 1 h post-exercise and returning to baseline at the 2 h time point (**Fig. 3f,g**). Urine urate levels were unchanged throughout the protocol (**Fig. 3h,i**). We conclude that urine Lac-Phe levels exhibit dynamic regulation following sprint exercise.

### Genetic ablation of SLC17 transporters reduces urine excretion of Lac-Phe

To directly test the *in vivo* contribution of SLC17 transporters to renal efflux of Lac-Phe in mice, we obtained knockout mice that are genetically deficient in either SLC17A1 or SLC17A3. SLC17A1 and A3 were selected because these two transporters exhibited the most robust Lac-Phe efflux activity in cells (**Fig. 2b**) and also high expression in the kidney (**Fig. 3a**). While both SLC17A1-KO and SLC17A3-KO mice were generated as part of the Knockout Mouse Project (KOMP) (22), the biochemical, molecular, and physiologic phenotypes of these animals have not been previously reported.

The SLC17A1/3 genes are localized in very close proximity (<20 kb) on mouse chromosome 13 (**Fig. 4a**). First, we characterized SLC17A1-KO mice, which were produced via CRISPR-Cas9 deletion of exon 3 (**Fig. 4a**). Complete loss of *SLC17A1* mRNA was observed from kidneys of SLC17A1-KO mice (**Fig. 4b**). The mRNA levels of the other SLC17 family members were not different, establishing specific deletion of only SLC17A1 in this genetic model (**Fig. 4b**). We were unable to robustly detect SLC17A1 protein by shotgun proteomics of whole kidney (**Table S2**). In SLC17A1-KO mice, a ∼30% reduction of urine Lac-Phe levels compared to WT littermates was observed (**Fig. 4c**); by contrast, the levels of other urine solutes, such as urate, were unchanged (**Fig. 4d**). In a separate cohort of mice, baseline plasma Lac-Phe were also unaltered in SLC17A1-KO mice (**Fig. 4e**).

**Fig. 4.**
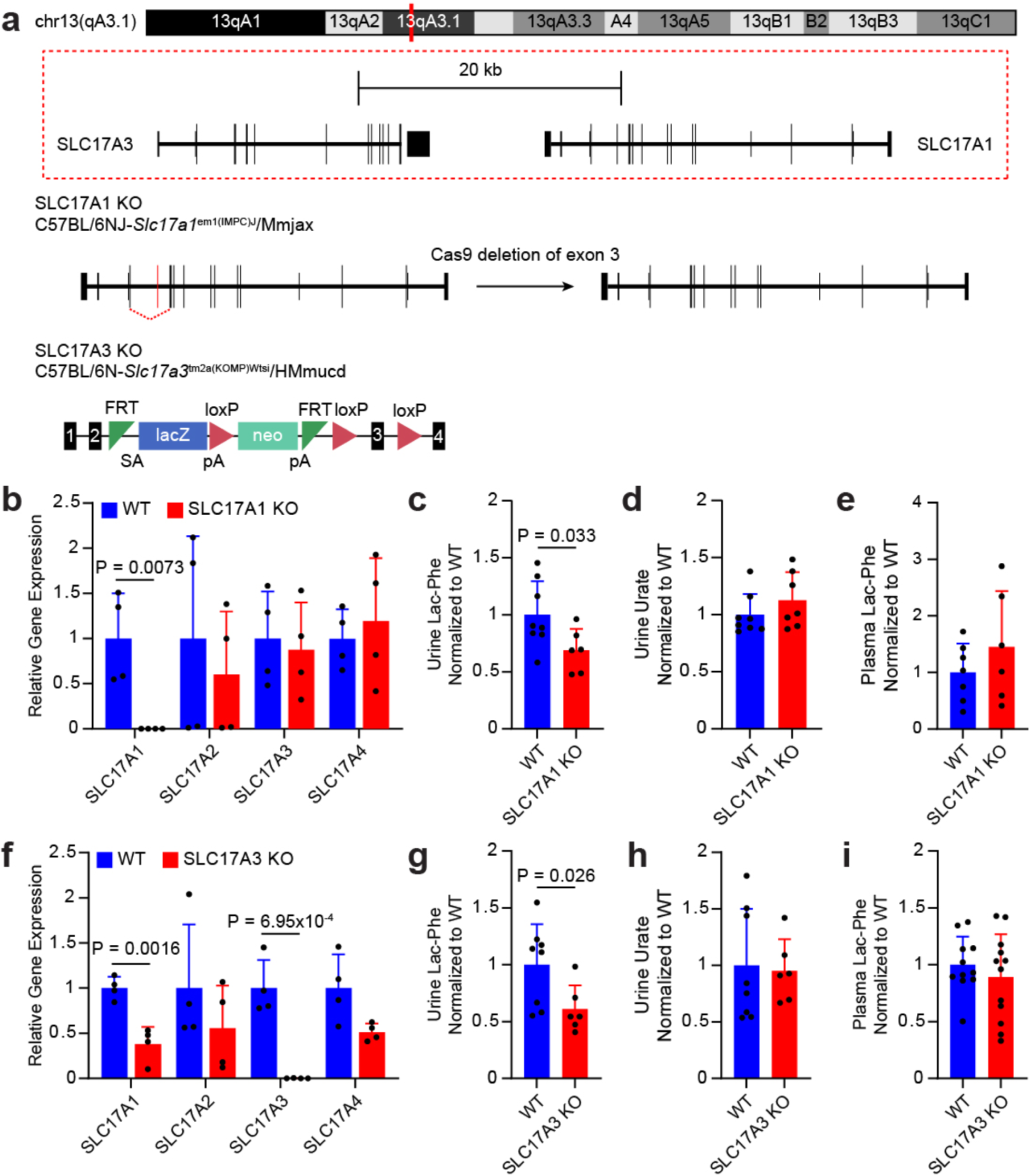
Reduced urine Lac-Phe levels in SLC17A1- and SLC17A3-KO mice. (**a**) Schematic of the SLC17A1 and SLC17A3 genes on mouse chromosome 13 and the genetic modifications resulting in the SLC17A1-KO or SLC17A3-KO mouse. (**b**) mRNA levels of the indicated genes from kidneys of WT or SLC17A1-KO mice, N=4/group. (**c,d**) Urine Lac-Phe (**c**) or urate (**d**) levels from male 3-5 month old WT or SLC17A1-KO mice. N=7-8/group. (**e**) Plasma Lac-Phe levels from 3-7 month old male and female WT or SLC17A1-KO mice, N=6-7/group. (**f**) mRNA levels of the indicated genes from kidneys of WT or SLC17A3-KO mice, N=4/group. (**g,h**) Urine Lac-Phe (**g**) or urate (**h**) levels from 1-3 month old male and female WT or SLC17A3-KO mice, N=6-8/group. (**i**) Plasma Lac-Phe levels from 2-3 month old male WT or SLC17A1-KO mice, N=6-7/group. Data are shown as mean ± SD. P-values for (b) and (f) were calculated using multiple *t*-tests with a false discovery rate approach of two-stage step-up method of Benjamini, Krieger and Yekutieli. P-values for (c-e) and (g-i) were calculated by Student’s two-sided *t*-test.

Next, we examined SLC17A3-KO mice. These animals harbor the so called tm2a KO first allele in which a splice acceptor and polyadenylation sequence are inserted in between exons 2 and 3. In kidneys from the SLC17A3-KO mice, *Slc17a3* mRNA was completely absent (**Fig. 4f**). We also observed a ∼60% partial reduction of *Slc17a1* mRNA and no changes in *Slc17a2* or *3* mRNA in this model (**Fig. 4f**). Shotgun proteomics revealed complete loss of SLC17A3 protein in knockout mice (**Table S3**). The partial reduction of *Slc17a1* mRNA may reflect complex and long-range interactions of the tm2a allele modification in the *Slc17a3* gene. In urine, Lac-Phe levels were reduced by 40% in SLC17A3-KO mice compared to WT controls (**Fig. 4g**) without any changes in urine urate levels (**Fig. 4h**). In a separate cohort of SLC17A3-KO mice, baseline plasma Lac-Phe levels were also not different (**Fig. 4i**). Our efforts to obtain double SLC17A1/3-KO mice via breeding were unsuccessful, which likely reflects the close physical proximity of these two alleles and consequently lack of independent recombination. We conclude that SLC17A1 and SLC17A3 are physiologic regulators of urine, but not plasma Lac-Phe levels.

After a single bout of treadmill exercise, mRNA levels of *Slc17a2* was reduced in the kidney, whereas mRNA levels of *Slc17a1*, *Slc17a3*, and *Slc17a4* were unchanged (**Fig. S8a**). We were unable to collect urine after treadmill exercise in mice, likely because the mice had already voided their bladders during the treadmill activity. Immediately after a single bout of exhaustive treadmill exercise, plasma Lac-Phe levels were still not different in either SLC17A1-KO or SLC17A3-KO mice (**Fig. S8b,c**).

Both SLC17A1-KO and SLC17A3-KO mice had normal kidney histology as assessed by hematoxylin and eosin (H&E) staining (**Fig. S9a,b**). In addition, plasma markers of renal function were unchanged in both genotypes (**Table S4** and **Table S5**). A blood metabolic panel also did not reveal any differences in either SLC17A1-KO and SLC17A3-KO mice (**Table S4** and **Table S5**). Body weights and food intake were also not different in SLC17A1-KO and SLC17A3-KO mice compared to WT controls on high fat diet (**Fig. S9c,d**), consistent with an absence of changes in circulating Lac-Phe levels in these animals. Lastly, we subjected a cohort of SLC17A3-KO and WT mice to a combined high fat diet/treadmill exercise protocol, but also did not detect any genotype-dependent effects on body weight or food intake (**Fig. S9e**).

After a single administration of Lac-Phe (50 mg/kg, IP), SLC17A3-KO mice had reduced Lac-Phe levels in urine compared to WT mice (**Fig. S10a**). We did not detect any differences in urine Lac-Phe in WT versus CNDP2-KO mice (**Fig. S10b**).

## Discussion

Here we provide several independent lines of evidence to show that SLC17 family members are physiologic urine Lac-Phe transporters: (1) overexpression of SLC17A1 and SLC17A3 dramatically increases Lac-Phe levels in conditioned media of cultured cells, a biochemical effect that is synergistic with the Lac-Phe biosynthetic enzyme CNDP2; (2) individual genetic ablation of SLC17A1 or SLC17A3 in mice reduces urine Lac-Phe levels; and (3) urine Lac-Phe levels exhibit strong genetic associations with the *SLC17A1-4* locus in humans. These data establish the first physiologic plasma membrane transporters for Lac-Phe *in vivo* and uncover a molecular mechanism involved in the renal excretion of this signaling metabolite.

Previously, SLC17 family members have been characterized to be plasma membrane transporters involved in renal excretion of urate. Supporting this hypothesis, SLC17 transporters exhibit urate transport activity *in vitro* (18) and polymorphisms in the *SLC17* locus are genetically associated with plasma urate levels in humans (23). Our studies further build on these observations by demonstrating that Lac-Phe is also an endogenous substrate of the SLC17 transporters. In addition, our untargeted metabolomics of cells transfected with SLC17 family members, as well as recent genome-wide association studies in humans, would also suggest that there are potentially many more physiologic SLC17 substrates beyond these alone (15). The ability to form anions, which is a characteristic present in both Lac-Phe and other annotated SLC17 substrates (e.g., urate, p-aminohippuric acid, acetylsalicylic acid), appears to be a common feature shared amongst SLC17 substrates.

Our mouse knockout studies demonstrate that each SLC17A1 and SLC17A3 contribute to renal excretion of Lac-Phe. Notably, that only partial deficiency of urine Lac-Phe is observed in each SLC17A1- and SLC17A3-KO mice is consistent with functional redundancy between these two solute carriers. We did not observe changes in urine urate levels with individual genetic ablation of SLC17A1/3 which may be due to a contribution of ABCG2 as a urate transporter (24) and/or the presence of uricase in mice which could catabolize excess urate to allantoin (25). In the future, the generation of double SLC17A1/3-KO mice, or potentially mice deficient in all four SLC17 transporters, may clarify whether there are additional, non-SLC17-dependent mechanisms for regulating urine Lac-Phe levels. In addition, kidney-specific SLC17A1 or SLC17A3 knockout mice, once available, could also help to further refine the importance of the renal expression of these transporters. Potent, selective, and in vivo-active SLC17A1/3 inhibitors, once developed, may also enable pharmacological interrogation of this pathway. Beyond Lac-Phe, in humans many additional urine metabolites have been associated with the *SLC17A1-4* locus, suggesting that these transporters have a large number of endogenous physiologic substrates.

The key insight that enabled identification of SLC17 carriers as Lac-Phe transporters originated from correct determination of the chemical structure of “X-15497/1-carboxyethylphenylalanine.” The similarity in MS/MS between Lac-Phe and 1-carboxyethylphenylalanine, especially in the m/z = 88 daughter ion, likely contributed to the initially incorrect structure. This correction underscores the importance of generating chemical standards and using differential retention time for high confidence structural assignment of metabolites. Our studies also enable the correct reinterpretation of “1-carboxyethylphenylalanine” in other public metabolomic datasets as Lac-Phe (26).

While SLC17A1 and SLC17A3 transporters clearly mediated Lac-Phe excretion to urine, we detected no effects on plasma Lac-Phe levels in the SLC17A1- or SLC17A3-KO mice. Therefore, the SLC17A1/3 transporters are not involved in Lac-Phe efflux to blood plasma, nor do they substantially contribute to clearance of plasma Lac-Phe levels. One explanation for this observation is that SLC17A1/3 transporters are localized to the apical plasma membrane and face the urine, and consequently only influence Lac-Phe levels in this matrix (27, 28). Future work will be required to uncover the molecular identity of the Lac-Phe transporter that mediates its efflux from its sites of production (e.g., Cndp2+ cells) to blood plasma. It seems plausible that different cell types might use different Lac-Phe transporters to efflux Lac-Phe into different body fluids. For instance, while ABCC5-KO mice do not have any differences in plasma Lac-Phe levels (9), it may be possible that ABCC5 physiologically contributes to Lac-Phe efflux in another fluid (e.g., urine, cerebral spinal fluid). Whether Lac-Phe can be transported in or out of cells lacking CNDP2 and/or SLC17A1/3 also remains an important question for the future. In addition, while SLC17-dependent excretion of Lac-Phe is indeed a physiologic process, this pathway does not appear to be a major pathway for the clearance of Lac-Phe from the circulation. Additional biochemical pathways for Lac-Phe regulation in the circulation, such as its potential uptake and degradation from blood plasma, remain to be defined.

## Acknowledgements

We thank members of the Long lab for helpful discussions. This work was supported by the NIH (DK124265 and DK136526 to JZL; GM113854 to VLL), the Stanford Diabetes Research Center (P30DK116074), and the Wu Tsai Human Performance Alliance (fellowship to VLL and ALW and research grant to JZL), and the Phil & Penny Knight Initiative for Brain Resilience at the Wu Tsai Neurosciences Institute (research grant to J.Z.L.). The work of PS was supported by the German Research Foundation (DFG) Project-ID 523737608 (SCHL 2292/2-1). The work of NS and AK was supported by German Research Foundation (DFG) project ID 431984000 (SFB 1453).

## Data availability

All data generated or analyzed during this study are included within the published articles and its supplementary information.

## Methods

### Cell line culture

All cell lines were obtained from the American Type Culture Collection (ATCC) and grown in an incubator at 37°C with 5% CO2. HEK293T and RAW264.7 cells were maintained in DMEM with 10% FBS and 2% penicillin-streptomycin. TKPTS cells were maintained in DMEM F-12 with 10% FBS and 2% penicillin-streptomycin.

### General animal information

Animal experiments were conducted according to the Stanford University Administrative Panel on Laboratory Animal Care approved protocols. Mice were maintained in 12-h light-dark cycles at 22°C and 50% relative humidity. Unless otherwise indicated, mice were fed a standard irradiated rodent chow diet. The high-fat diet used was from Research Diets (D12492, 60% kcal from fat). SLC17A1 animals were obtained from the International Mouse Phenotyping Consortium (IMPC, C57BL/6NJ-Slc17a1em1(IMPC)J/Mmjax) and SLC17A3 animals were obtained from the Knockout Mouse Project (KOMP, C57BL/6N-Slc17a3tm2a(KOMP)Wtsi/HMmucd, 049670-UCD). Sample sizes were determined on the basis of previous experiments using similar methodologies.

### Chemicals

N-Lactoyl Phenylalanine (catalogue # 318943) was purchased from Novo Pro Labs and 1-Carboxyethylphenylalanine (ENAH95E73EBB) was purchased from Sigma Aldrich.

### *In vitro* transporter secretion assays

HEK293T cells were plated in 10 cm plates at 5 million cells/plate and the next day transiently transfected with the indicated plasmids using PolyFect (Qiagen 301105) according to the manufacturer’s instructions. RAW 264.7 cells were transfected with Lipofectamine LTX reagent with PLUS reagent according to manufacturer’s instructions. One-day post-transfection, cells were re-plated into 12-well plates at 70-80% confluence. The next day cells were put into 0.5 ml of serum free media. After overnight incubation, 400 μl of media was removed and 20 μl of 1 M hydrochloride added to acidify the media and protonate Lac-Phe. 400 μl of ethyl acetate was added into each sample and vortexed for 30 seconds to extract Lac-Phe into the organic layer. 300 μl from the top layer was transferred to a new Eppendorf tube and dried down under a stream of nitrogen. The residue was re-suspended in 100 ul of an 80:20 mixture of acetonitrile/water. The mixture was centrifuged at 4 °C for 10 minutes at 15,000 rpm and the supernatant was transferred to a LC-MS vial. For collection of intracellular metabolites, cells were wash 2 times with PBS and lysed in 150 ul of a 2:1:1 acetonitrile:methanol:water mixture. The cells were scraped from the well and both cell lysate and supernatant were transferred to an Eppendorf tube. The samples were then centrifuged at 4 °C for 10 minutes at 15,000 rpm and the supernatant was transferred to a LC-MS vial.

### Untargeted measurements of metabolites by LC-MS

Untargeted metabolomics measurements were performed on an Agilent 6520 Quadrupole Time-of-Flight (Q-TOF) LC/MS. Mass spectrometry analysis was performed using electrospray ionization (ESI) in negative mode. The dual ESI source parameters were set as follows, the gas temperature was set at 250 °C with a drying gas flow of 12 l/min and the nebulizer pressure at 20 psi. The capillary voltage was set to 3500 V and the fragmentor voltage set to 100 V. Separation of polar metabolites was conducted on a Luna 5 μm NH2 100 Å LC column (Phenomenex 00B4378-E0) with normal phase chromatography. Mobile phases were as follows: Buffer A, 95:5 water/acetonitrile with 0.2% ammonium hydroxide and 10 mM ammonium acetate. Buffer B, acetonitrile. The LC gradient started at 100% B with a flow rate of 0.2 ml/min from 0-2 min. The gradient was then increased linearly to 50% A/50% B at a flow rate of 0.7 ml/min from 2-20 minutes. From 20-25 minutes the gradient was maintained at 50% A/50% B at a flow rate of 0.7 ml/min.

### Generation of SLC17A1 KO TKPTS Cells

The plentiCRISPRv2 system developed by the Zhang lab was used to generate the SLC17A1-KO TKPTS cell line. The sgRNA used was 5’-TGGGCACCTCCCTTAGAACG-3’.

Following Zhang lab protocols, oligonucleotides for the sgRNA and reverse complement sequences were synthesized and cloned into the plentiCRISPRv2 vector. Lentivirus particles were generated in the HEK293T cell line using PEI for the co-transfection of the cloned plentiCRISPRv2 plasmid with the viral packing psPAX2 plasmid, and viral envelope pMD2.G plasmid. A non-cutting plentiCRISPRv2 plasmid was used as a control. The lentivirus was harvested after 48 hours and filtered through a 0.45 uM filter. The supernatant was then mixed in a 1 to 1 ratio with polybrene to a final concentration of 8 ug/ml polybrene. This mixture was added to TKPTS cells for transduction. Two days later, the cells were split into a 10 cm plate and underwent selection with 5 ug/ml puromycin for 3-6 days.

### Human exercise study

10 healthy individuals, 6 male, 4 female, age 28 ± 5 (mean ± SD), were enrolled and consented to participate in the exercise study approved by the Stanford University Institutional Review Board (IRB-65281). Participants arrived at the test facility and urine collected prior to exercise training. The participants then underwent a sprint exercise Wingate test. The Wingate test was conducted on an ergometer bike in which there was a 5 min warm-up followed by three bouts of 30 sec all-out sprint interspersed by 4 min of active recovery. Urine samples were collected before and 1 and 2 hours post-exercise for LC-MS analysis.

### Mouse running protocols

For mouse running studies a 6 lane Columbus Instruments animal treadmill (product 1055-SRM-D65) was used. Mice were first acclimated to the treadmill for 5 minutes before the running protocol began. For acute, intense running experiments, the treadmill speed begun at 7.5 m/min at a 4° incline. Every 3 minutes, the speed and incline were increased by 2.5 m/min and 2°, respectively until a maximum speed of 40 m/min and incline of 30° was reached. Mice were run on the treadmill until exhaustion which was defined as when mice remained on the shocker at the end of the treadmill for longer than 5 seconds. For chronic running experiments, mice were exercised 5 days/week (Monday-Friday). The mice were also kept on high fat diet (60% kcal from fat). The treadmill running was performed at a constant 5° incline and began at a speed of 6 m/min. Speed was increased by 2 m/min every 5 minutes until a maximum speed of 30 m/min. Mice were once again stopped upon reaching exhaustion and run times were normalized between the two genotypes.

### Preparation of plasma and urine samples for LC-MS analysis

Plasma was collected from mice via a submandibular bleed into lithium heparin tubes (BD, 365985) and kept on ice. The blood was centrifuged at 4°C at 5,000 r.p.m. for 5 min to separate out the top plasma layer which can be frozen at -80°C. To collect urine, mice were scruffed over an eppendorf tube and gently stroked until they urinated; samples were also stored at -80°C. To extract polar metabolites for LC-MS analysis, 150 ul of a 2:1 mixture of acetonitrile:methanol was added to 50 ul of either plasma or urine. The mixture was centrifuged at 4°C at 15,000 r.p.m. for 10 min and the supernatant was transferred to a LC-MS vial.

### Breeding and genotyping of SLC17A1 and SLC17A3 KO mice

SLC17A1-KO and WT animals as well as SLC17A3-KO and WT animals were generated by heterozygous breeding crosses. Genotyping was performed as follows: Tail clippings were obtained from littermates and boiled for 30 minutes at 95 °C in 100 μl of 50 mM NaOH to extract genomic DNA. The solution was neutralized by adding 21 μl of 0.5 M Tris (pH 7.2). For SLC17A1 animals, PCR reactions were performed by using primers for either the *SLC17A1* WT allele (forward, 5’-CAGTTCCAGGGTTCTGTTCC-3’; reverse, 5’-GGTGACCCTGTGTTGTTCAC-3’) or SLC17A1 mutant allele (forward, 5’-TGTGCAACTGTCAACCAGAGT-3’; reverse, 5’-CCAGTCTAGTGAGTGCTGTCGTC-3’). The Promega GoTaq master mix was used for the PCR reaction. Each 25 μl reaction consisted of 12.5 μl of the promega master mix (M7122), 2.5 μl of a 10 μM mixture of forward and reverse primers, 2 μl of genomic DNA, and 8 μl of ultrapure water. The thermocycling program on BioRad C1000 Touch Thermo Cycler began with an initial 1 min 30 seconds at 98°C, followed by cycles of 30 seconds at 98°C, 30 seconds at 60 °C, and 30 seconds at 72 °C (30 cycles), followed by 5 minutes at 72 °C and finally held at 4 °C. Samples were run on a 2% agarose gel with 0.2 mg/ml EtBr. WT alleles are expected to yield a PCR product 130 base pairs in size while KO alleles are expected to yield PCR products that are 150 base pairs in size. For SLC17A3 animals, PCR reactions were performed by using primers for either the *SLC17A3* WT allele (forward, 5’-TCAAATGAAAGTTTATCCTCAATGCCTG-3’; reverse, 5’-AGATATATAGAGCCTGTGTCCACTGG-3’) or the PreCre allele (forward, 5’-GGGATCTCATGCTGGAGTTCTTCG-3’; reverse, 5’-AGATATATAGAGCCTGTGTCCACTGG-3’). The Promega GoTaq master mix was used for the PCR reaction. Each 15 μl reaction consisted of 7.5 μl of the promega master mix (M7122), 0.45 μl of a 20 μM mixture of forward and reverse primers, 1 μl of genomic DNA, and 6.05 μl of ultrapure water. The thermocycling program on BioRad C1000 Touch Thermo Cycler began with 5 min at 94 °C, followed by cycles of 15 seconds at 94 °C, 30 seconds at 65 °C (-1 °C/cycle), 40 seconds at 72 °C (10 cycles), followed by 15 seconds at 94 °C, 30 seconds at 55°C, 40 seconds at 72 °C (30 cycles), and ending with 5 minutes at 72 °C and finally held at 4 °C. Samples were run on a 2% agarose gel with 0.2 mg/ml EtBr. WT alleles are expected to yield a PCR product 348 base pairs in size while KO alleles are expected to yield PCR products that are 559 base pairs in size.

### Shotgun proteomics

Samples were reduced with 10 mM dithiothreitol (DTT) for 20 minutes at 55 degrees Celsius, cooled to room temperature and then alkylated with 30 mM acrylamide for 30 minutes. They were then acidified to a pH ∼1 with 2.6 ul of 27% phosphoric acid, dissolved in 165 uL of S-trap loading buffer (90% methanol/10% 1M triethylammonium bicarbonate (TEAB)) and loaded onto S-trap microcolumns (Protifi, C02-micro-80). After loading, the samples were washed sequentially with 150 ul increments of 90% methanol/10% 100mM TEAB, 90% methanol/10% 20 mM TEAB, and 90% methanol/10% 5 mM TEAB solutions, respectively. Samples were digested at 47 °C for two hours with 600 ng of mass spectrometry grade Trypsin/LysC mix (Promega, V5113). The digested peptides were then eluted with two 35 µl increments of 0.2% formic acid in water and two more 40uL increments of 80% acetonitrile with 0.2% formic acid in water. The four elutions were consolidated in 1.5 ml S-trap recovery tubes and dried via SpeedVac (Thermo Scientific, San Jose CA). Finally, the dried peptides were reconstituted in 2% acetonitrile with 0.1% formic acid in water for LC-MS analysis.

Mass spectrometry experiments were performed using an Orbitrap Exploris 480 mass spectrometer (Thermo Scientific, San Jose, CA) attached to an Acquity M-Class UPLC system (Waters Corporation, Milford, MA). The UPLC system was set to a flow rate of 300 nl/min, where mobile phase A was 0.2% formic acid in water and mobile phase B was 0.2% formic acid in acetonitrile. The analytical column was prepared in-house with an I.D. of 100 microns pulled to a nanospray emitter using a P2000 laser puller (Sutter Instrument, Novato, CA). The column was packed with Dr. Maisch 1.9 micron C18 stationary phase to a length of approximately 25 cm. Peptides were directly injected onto the column with a gradient of 3-45% mobile phase B, followed by a high-B wash over a total of 80 minutes. The mass spectrometer was operated in a data-dependent mode using HCD fragmentation for MS/MS spectra generation.

RAW data were analyzed using Byonic v4.4.1 (Protein Metrics, Cupertino, CA) to identify peptides and infer proteins. A concatenated FASTA file containing Uniprot Mus musculus proteins and other likely contaminants and impurities was used to generate an *in silico* peptide library. Proteolysis with Trypsin/LysC was assumed to be semi-specific allowing for N-ragged cleavage with up to two missed cleavage sites. Both precursor and fragment mass accuracies were held within 12 ppm. Cysteine modified with propionamide was set as a fixed modification in the search. Variable modifications included oxidation on methionine, histidine and tryptophan, dioxidation on methionine and tryptophan, deamidation on glutamine and asparagine, and acetylation on protein N-terminus. Proteins were held to a false discovery rate of 1% using standard reverse-decoy technique.

### Statistics

All data are expressed as mean ± SD or SEM as indicated in the legends. A student’s t-test was used for pair-wise comparisons. For comparisons of multiple groups, P-values were calculated by one-way ANOVA with Dunnett T3 post-hoc tests. Unless otherwise specified, statistical significance was set as P < 0.05.

## Author contributions

Conceptualization: VLL, JZL; investigation: VLL, SX, PS, NS, ALW, JS, ASHT; writing – original draft, VLL, JZL; writing – reviewing & editing, VLL, PS, NS, AK, JZL; resources EDK, AK, JZL, supervision and funding acquisition, AK, JZL.

## Competing interests

The authors declare no competing interests.

## Notes

### Competing Interest Statement

The authors have declared no competing interest.

