## Supplemental Information for "SLC17 transporters mediate renal excretion of Lac-Phe in mice and humans"

Li et al.

Contents

**Fig. S1-S10 and legends**

**Table S1-5 legends**

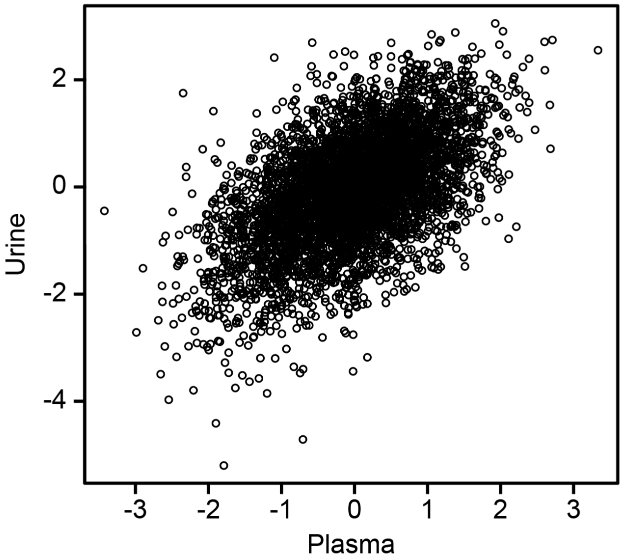


**Fig. S1. Correlation of urine and plasma Lac-Phe levels.**
Re-analysis of paired urine and plasma Lac-Phe levels from Schlosser P, et al. (2023) Genetic studies of paired metabolomes reveal enzymatic and transport processes at the interface of plasma and urine. *Nat Genet* 55(6):995–1008.

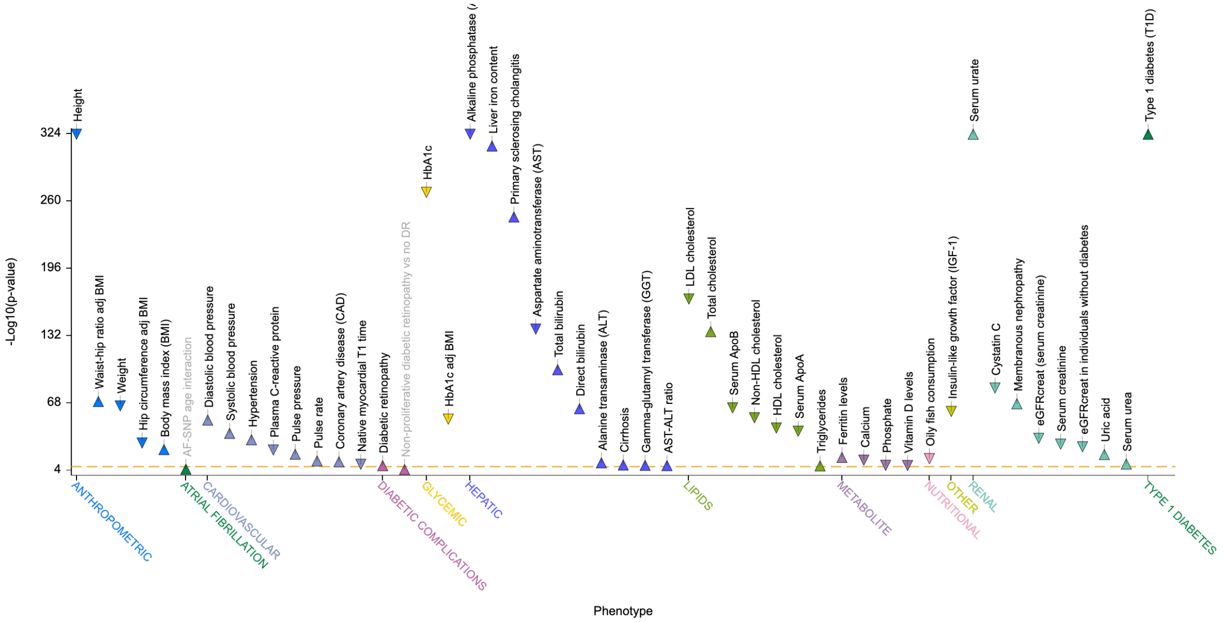


**Fig. S2. Phenotype associations of the human *SLC17* locus.**

Most significant variant associations in the region 6:25,733,143-25,882,280 (Ancestry: All) from the Type 2 Diabetes Knowledge Portal. Associations are clumped by linkage disequilibrium. LD clumping was performed using PLINK (https://zzz.bwh.harvard.edu/plink/clump.shtml) with the following parameters: PLINK_P1 = 5e-8, PLINK_P2 = 1e-2, PLINK_R2 = 0.2, PLINK_KB = 250.

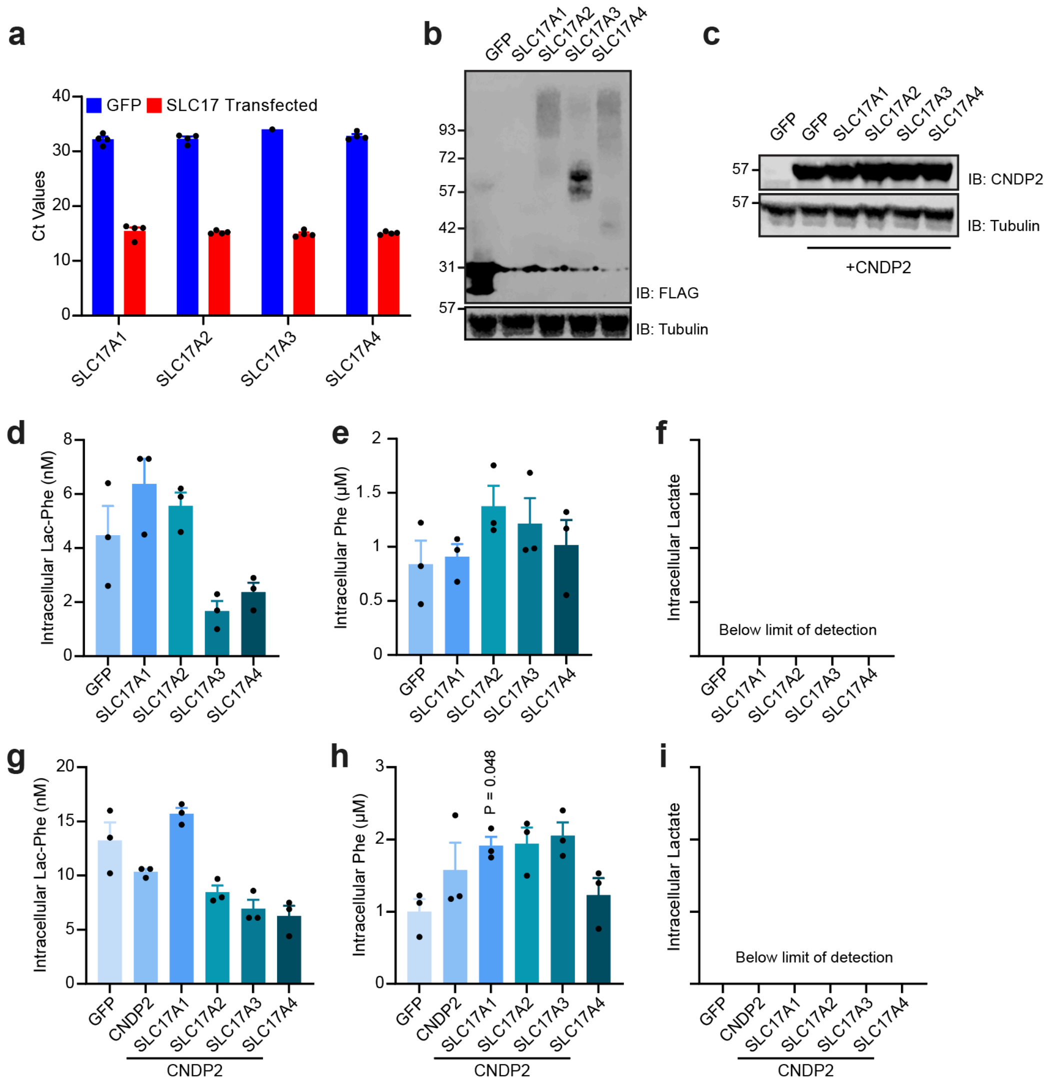


**Fig. S3. Additional characterization of SLC17-mediated Lac-Phe efflux in HEK293T cells.**
(a,b) Ct values by qPCR (a) and Western blot using an anti-flag (b, top) or anti-tubulin (b, bottom) antibody of HEK293T cells two days after transfection with the indicated cDNA. All SLC17 transporters contain a C-terminal flag tag.
(c) Western blot using an anti-CNDP2 (top) or anti-tubulin (bottom) antibody of HEK293T cells two days after co-transfection with the indicated plasmids and CNDP2.
(d-i) Intracellular concentrations of the indicated metabolites in HEK293T cells two days after transfection with the indicated cDNAs.
For (a), N=4/group. For (d-h), N=3/group. Data are shown as means ± SEM. P-values for (d-h) were calculated by one-way ANOVA with Dunnett T3 post-hoc tests.

Mouse RAW264.7 macrophages

Mouse TKPTS kidney proximal tubule cells

**a**

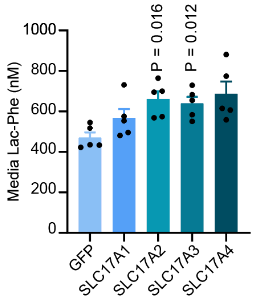


**b**

**Ct Values**

| | | | |
|---|---|---|---|
| **TBP** | 22.02 | 22.14 | 22.16 |
| **SLC17A1** | 25.95 | 26.26 | 26.36 |
| **SLC17A2** | 31.01 | 31.97 | 32.11 |
| **SLC17A3** | 33.46 | 32.63 | 34.30 |
| **SLC17A4** | 31.62 | 32.50 | 33.51 |

**c**

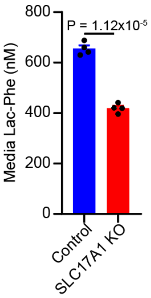


**Fig S4. Effect of SLC17 transporters on media Lac-Phe levels in additional cell types.**
(a) Media Lac-Phe concentrations from mouse macrophage RAW264.7 cells two days after transfection with the indicated cDNAs.
(b) Ct values of the indicated mRNAs from the mouse renal proximal tubule cell line TKPTS.
(c) Media Lac-Phe concentrations in control and SLC17A1-KO TKPTS cells.
For (a), N=5/group. For (c), N=4/group. Data are shown as means ± SEM. P-values for (a) were calculated by one-way ANOVA with Dunnett T3 post-hoc tests. P-value for (c) was calculated using a Student’s two-sided *t*-test.

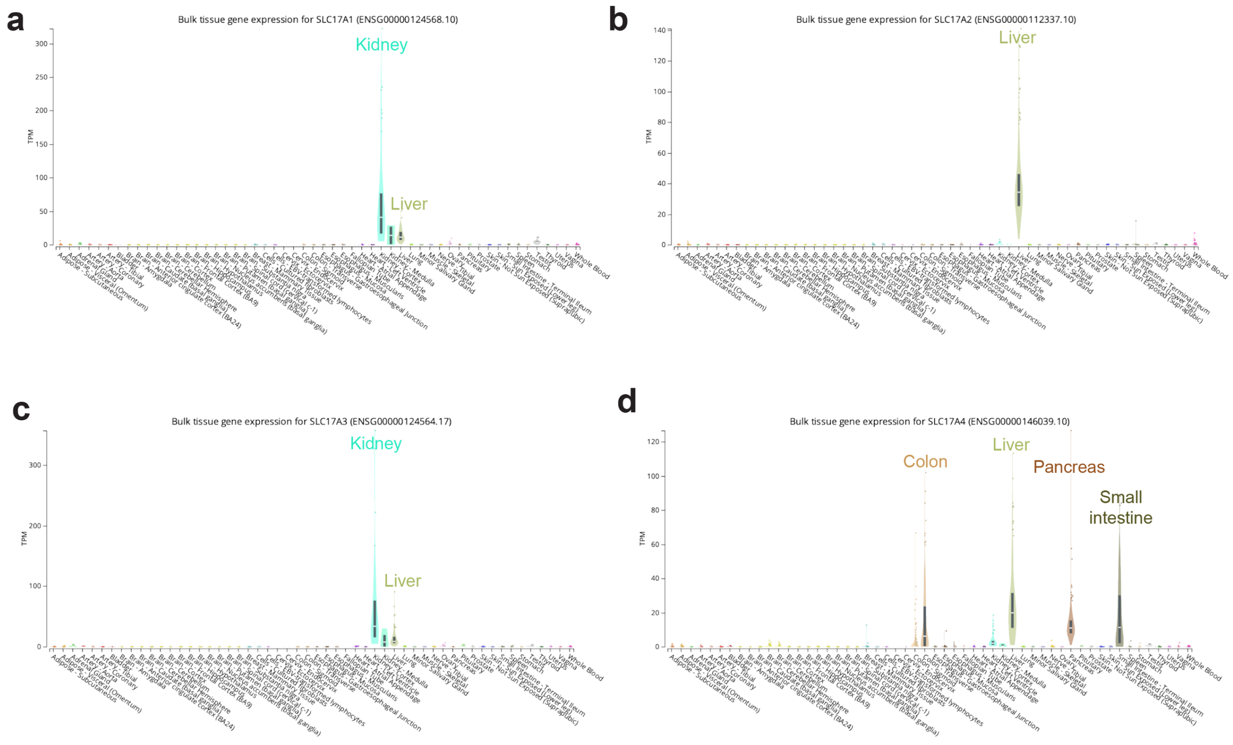


**Fig. S5. Expression of *SLC17* transporters in humans.**
(a-d) mRNA levels across tissues for human *SLC17A1* (a), *SLC17A2* (b), *SLC17A3* (c), and *SLC17A4* (d). Data were obtained from the GTEx Portal.

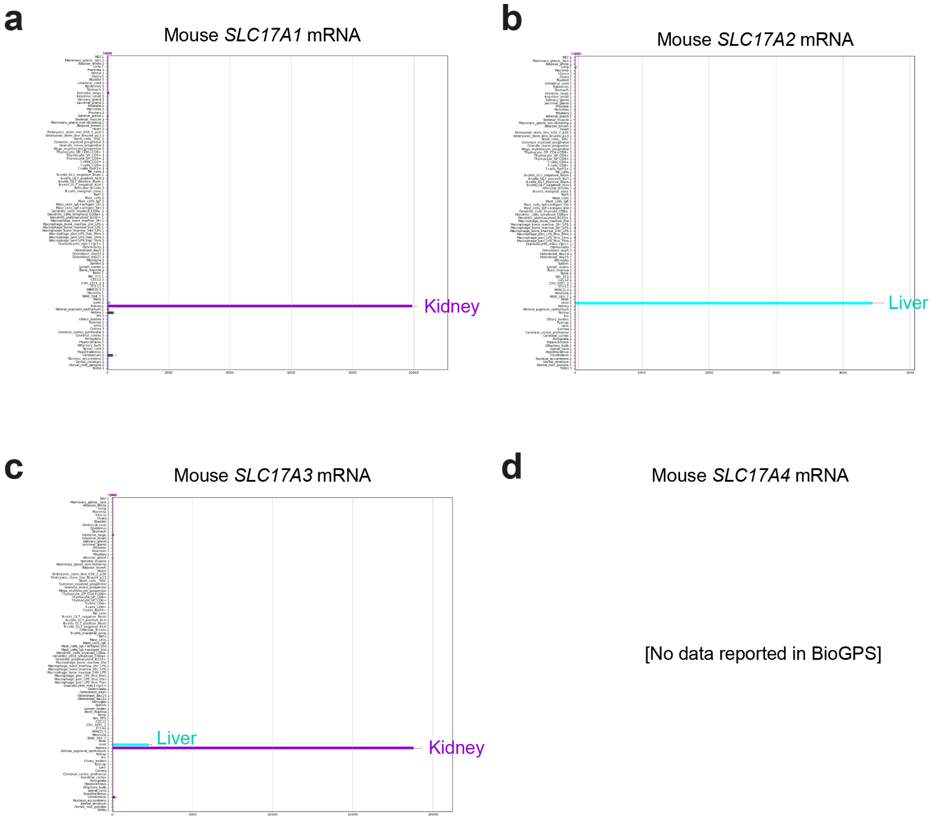


**Fig. S6. Expression of *Slc17* transporters in mice.**
(a-d) mRNA levels across tissues for mouse *Slc17a1* (a), *Slc17a2* (b), *Slc17a3* (c), and *Slc17a4* (d). Data were obtained from BioGPS.

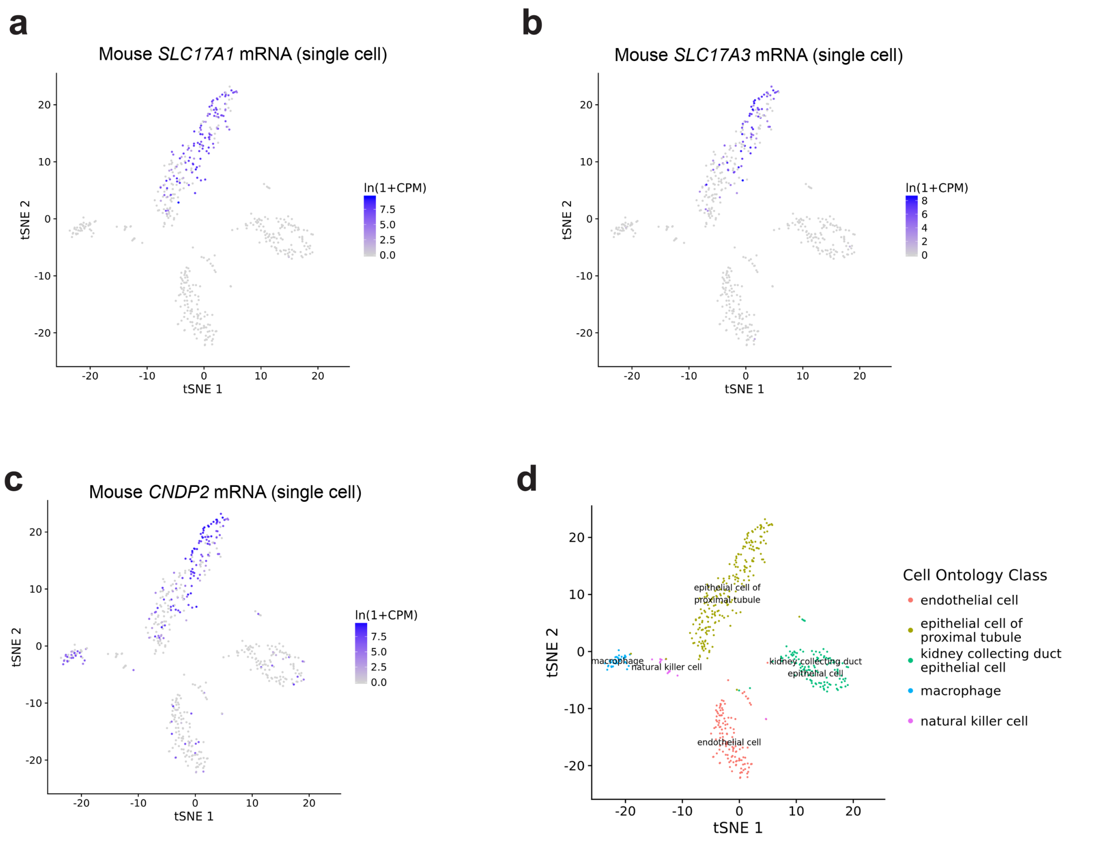


**Fig. S7. Single-cell mRNA expression data for *Slc17a1*, *Slc17a3,* and *Cndp2* in mice.** (a-d) Expression of mouse *Slc17a1* (a), *Slc17a3* (b), and *Cndp2* (c), and cell type annotation (d), of mouse kidney single cell RNAseq data obtained from Tabula Muris.

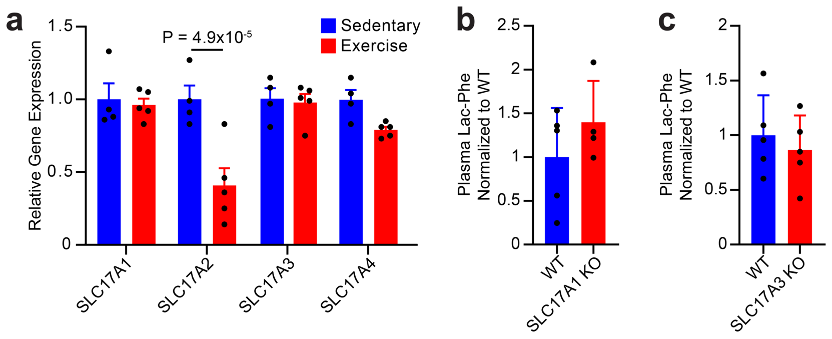


**Fig. S8. Additional characterization of kidney and blood following acute treadmill exercise in mice.**

(a) mRNA levels of the indicated genes in sedentary male 7-week old mice mice or after a single bout of exhaustive treadmill running (exercise).

(b, c) Plasma Lac-Phe levels in 3-4 month old male SLC17A1-KO (b) or 4-month old male SLC17A3-KO (c) mice after a single bout of exhaustive treadmill running.

For (a), N=5/group. For (b), N=4-5/group. For (c) N=5/group. Data are shown as mean ± SEM for (a) and as mean ± SD for (b,c). P-values were calculated by Student's *t*-test.

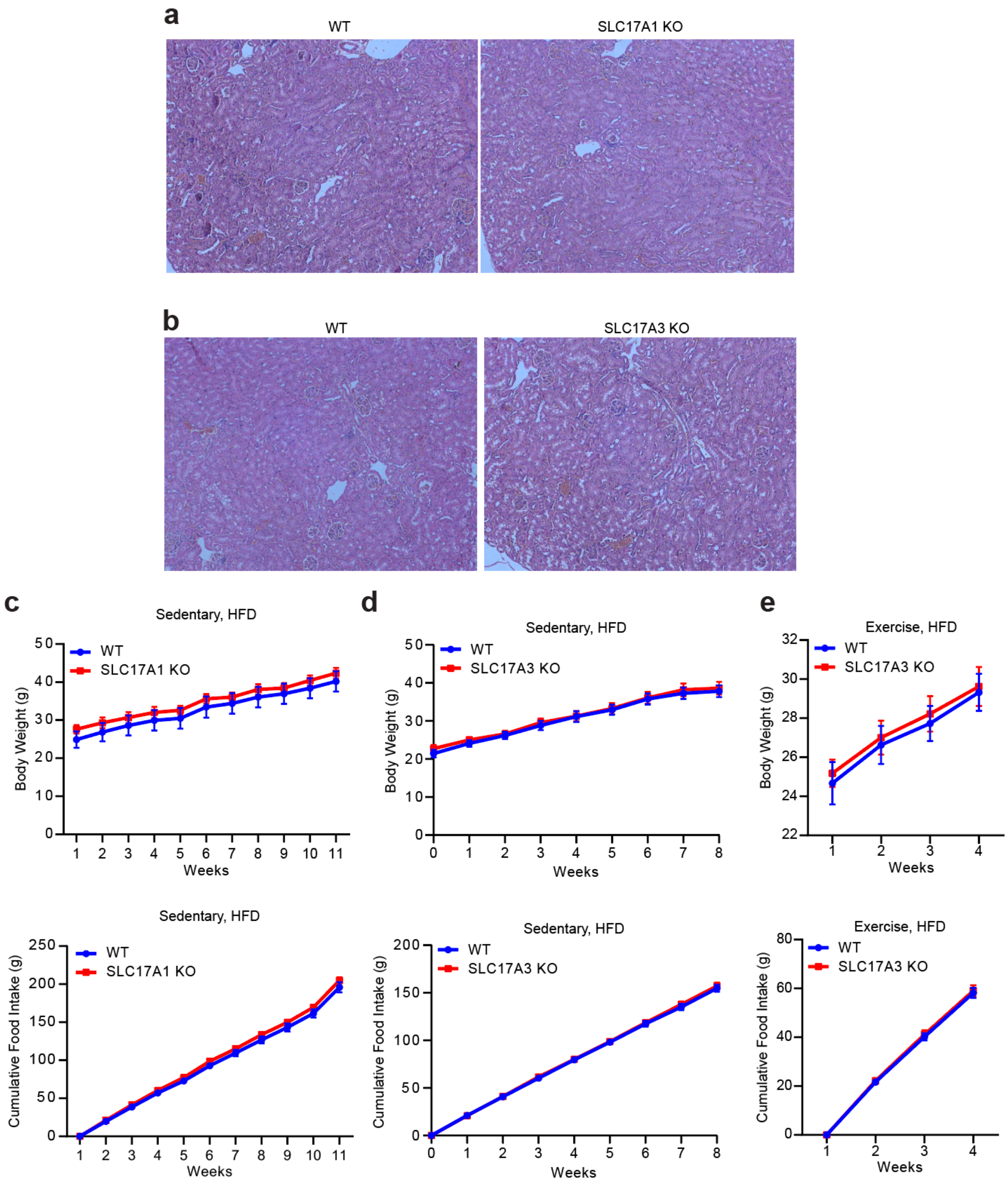


**Fig. S9. Additional characterization of SLC17A1- and SLC17A3-KO mice.**
(a,b) Representative H&E staining of kidney sections from 8-9 month old male WT and SLC17A1-KO (a) or SLC17A3-KO (b) mice.
(c-e) Body weights (top) and cumulative food intake (bottom) of high fat diet-fed sedentary 4-6 month old male SLC17A1-KO mice (c), 2-3 month old male SLC17A3-KO mice (d), or high fat diet-fed and exercised 3-4 month old male SLC17A3-KO mice (e). See **Methods** for the chronic exercise protocol.
For (c-e), data are shown as means ± SEM. For (c), N=8/group. For (d), N=11-12/group. For (e), N=8-9/group.

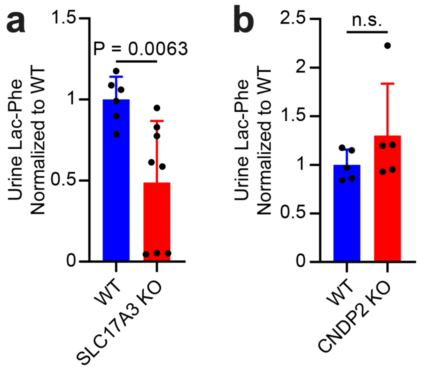


**Fig. S10. Additional characterization of urine Lac-Phe levels.**
(a) Levels of Lac-Phe in urine from 6-9 month old, mixed male and female WT or SLC17A3-KO mice 30 minutes after a single administration of Lac-Phe (50 mg/kg, IP), N=6-8/group.
(b) Levels of Lac-Phe in urine from 2-4 month old, mixed male and female WT or CNDP2-KO mice, N=5/group.
Data are shown as means ± SD. P-values were calculated by Student's *t*-test.

### Supplementary Table Legends

**Table S1.** Differential metabolomics by XCMS from media of HEK293T cells two days after transfection with GFP, SLC17A1, or SLC17A3.

**Table S2.** Shotgun proteomics of kidney from WT or SLC17A1-KO mice.

**Table S3.** Shotgun proteomics of kidney from WT or SLC17A3-KO mice.

**Table S4.** Blood lipid and renal panels in WT and SLC17A1-KO mice.

**Table S5.** Blood lipid and renal panels in WT and SLC17A3-KO mice.
